# Non-instrumental information has limited effects on bet size in risky decisions

**DOI:** 10.64898/2026.07.30.741906

**Authors:** Matt Jiwa, Dan Myles, Daniel Bennett

## Abstract

Recent research has suggested that the availability of non-instrumental information about the outcome of a risky choice increases risk appetite. In this study, we aimed to perform a conceptual replication of these findings and to examine the cognitive mechanisms underlying this effect. Across two experiments (N = 150, 102), we presented participants with mathematically fair gambles and allowed them to choose the size of their bet. Between trials, we varied the presence of non-instrumental information that would reveal the outcome ahead of time. In both experiments, we did not find consistent evidence for an effect of the availability of non-instrumental information on bet size. These findings suggest that the previously reported effects of non-instrumental information on risk appetite may have been an idiosyncratic feature of experimental design, rather than a more general phenomenon that characterises human decision making under risk.

---

The cognitive mechanisms underlying decisions made under risky or uncertain conditions have long been a cornerstone of cognitive science research. Early frameworks emphasised the role of expected utility (Von Neumann & Morgenstern, 1947), and were later expanded to incorporate subjective distortions (Tversky & Kahneman, 1992). These frameworks suggest that decisions made under risk should be based on the statistical properties of the possible outcomes therein. For example, when deciding whether to play a lottery or not, an agent should consider the likelihood of winning or losing as well as the associated payout or loss. Information that reduces uncertainty about these statistical properties can often be used to inform decision making in a way that affects the likelihood or magnitude of future rewards; for instance, clairvoyance of the winning lottery numbers could be used to select a winning ticket in advance. Information with these properties can be described as possessing “instrumental” value (Sharot & Sunstein, 2020). However, recent research has demonstrated that we may also value information that cannot be used to alter the probability or value of a future outcome (termed “non-instrumental” information; Bennett et al., 2016; Charpentier et al., 2018; Goh et al., 2021; Jiwa and Gottlieb, 2026; Van Lieshout et al., 2018). Importantly, unlike the fanciful clairvoyance example, signals containing non-instrumental information are a common feature of everyday gambling products: slot machine reel stops, lottery number draws, and scratch card panels are all examples of sequentially-revealed non-instrumental information that indicates a gamble’s outcome in advance of a final outcome, but that cannot be used to improve one’s odds of winning.

One recent study reported that the availability of this kind of non-instrumental information increases appetite for risk (Matthews et al., 2023). This study demonstrated that participants were more willing to place a bet using a stylised slot machine if the outcome was accurately signalled in advance by the progressive reveal of five slot windows. Computational modelling analyses revealed the effect of information on risk appetite was best described through its capacity to reveal the outcome of the lottery as soon as possible, resolving uncertainty early. However, the cognitive mechanisms underlying this effect remain unknown, and the task used by Matthews et al. also introduced a potential confound: in their task, non-instrumental information was revealed only if participants chose to put money into the slot machine (i.e., there was no way to view information without taking a risk on the slot machine). Consequently, decisions to play the slot machine may have reflected participants’ desire for non-instrumental information rather than inflation of their risk appetites by non-instrumental information.

In the present study, we aimed to address this potential confound and to examine the cognitive mechanisms underlying the effect of non-instrumental information availability on risk appetite. We first used a novel task in which participants chose how much money to risk in a fair gamble across trials that either rendered the outcome after a six-second delay, or signalled the outcome halfway through the delay period (thereby providing non-instrumental information). After failing to produce a strong conceptual replication of the original effect, we then altered the paradigm to increase its consistency with Matthews et al.’s task by adopting a five-stage sequential reveal of information while still decoupling demand for non-instrumental information from its effects on risk appetite.

## Methods

### Participants

Using the online participant recruitment platform Prolific (Palan & Schitter, 2018), we recruited samples of 150 participants in Experiment 1 (72 female, 77 male, 1 non-binary or gender diverse; *M* = 39.37, *SD* = 11.07) and 102 participants in Experiment 2 (50 female, 51 male, 1 non-binary or gender diverse; *M* = 37.76, *SD* = 11.09). Participation was restricted to those aged between 18 and 65, with fluent written and spoken English and no visual impairments. To ensure data was high-quality, we also applied a filter to recruit only participants with at least two prior Prolific submissions and with a minimum of 80% approval rate on those submissions. Each task took approximately 30 minutes to complete. For their participation, participants received 4 GBP, plus their winnings from the task (Experiment 1: *M* = 1.07 GBP, *SD* = 0.12 GBP; Experiment 2: *M* = 1.05 GBP, *SD* = 0.34 GBP). In both experiments, informed consent was provided by all participants, and research was conducted in accordance with the Declaration of Helsinki. All study protocols were approved by The University of Melbourne Human Research Ethics Committee (ID 33865).

### Procedure

All stimuli were presented using the jsPsych library (De Leeuw, 2015; version 7.3.1). Before commencing the main task, participants completed a short demographic survey to record their age, gender, education level, and past-year gambling participation. Respondents who had gambled in the past year were then prompted to complete the Problem Gambling Severity Index (PGSI; Ferris and Wynne, 2001). All participants were then presented with comprehensive written and visual instructions for the task, before completing 8 practice trials. Finally, they responded to a series of five simple true or false questions that checked their understanding of the task. If they responded incorrectly to any of these comprehension questions, they were taken back to the start of the instructions to review them, before returning to the understanding check questions with no additional practice trials.

Participants were instructed that they would start with 1000 grams of gold, and that they would receive a bonus payment at the end of the task based on the amount of gold they finished with at an exchange rate of 1 GBP per 1000 grams. At the beginning of each trial the participant was presented with an image of a “gold-refining machine” represented by a geometric box (Figure 1). Below each machine, we explicitly displayed the percentage probability that the machine would successfully increase the value of the gold it processed, as well as the potential return per gram of gold if the process was successful. This return was the reciprocal of the success probability: if the probability of successful refinement was 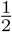, the machine would return 2 grams of gold for each gram put in, if it was 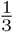, the machine would return 3 grams, and so on. Participants used a text box to indicate how many grams of ‘gold’ they wished to ‘refine’ (i.e., the bet amount), with responses restricted to integer values between 1 and 50. No time limit was imposed on these responses. There was then a six-second delay before a message was displayed on screen indicating the success or failure of the process and the value lost or gained. If refining was successful, the outcome was the product of the bet amount and the return per gram. If refining was unsuccessful, participants lost the bet amount. During the delay period the image of the machine remained on screen alongside the text “Refining in progress… “. The total amount of gold the participant possessed was then updated and displayed at the top of the screen at the start of the next trial. Bet amount was taken as an indicator of risk appetite, because placing larger bets increases the magnitude of both prospective gains in the case of a success and prospective losses in the case of a failure (cf. Demaree et al., 2008).

**Figure 1:**
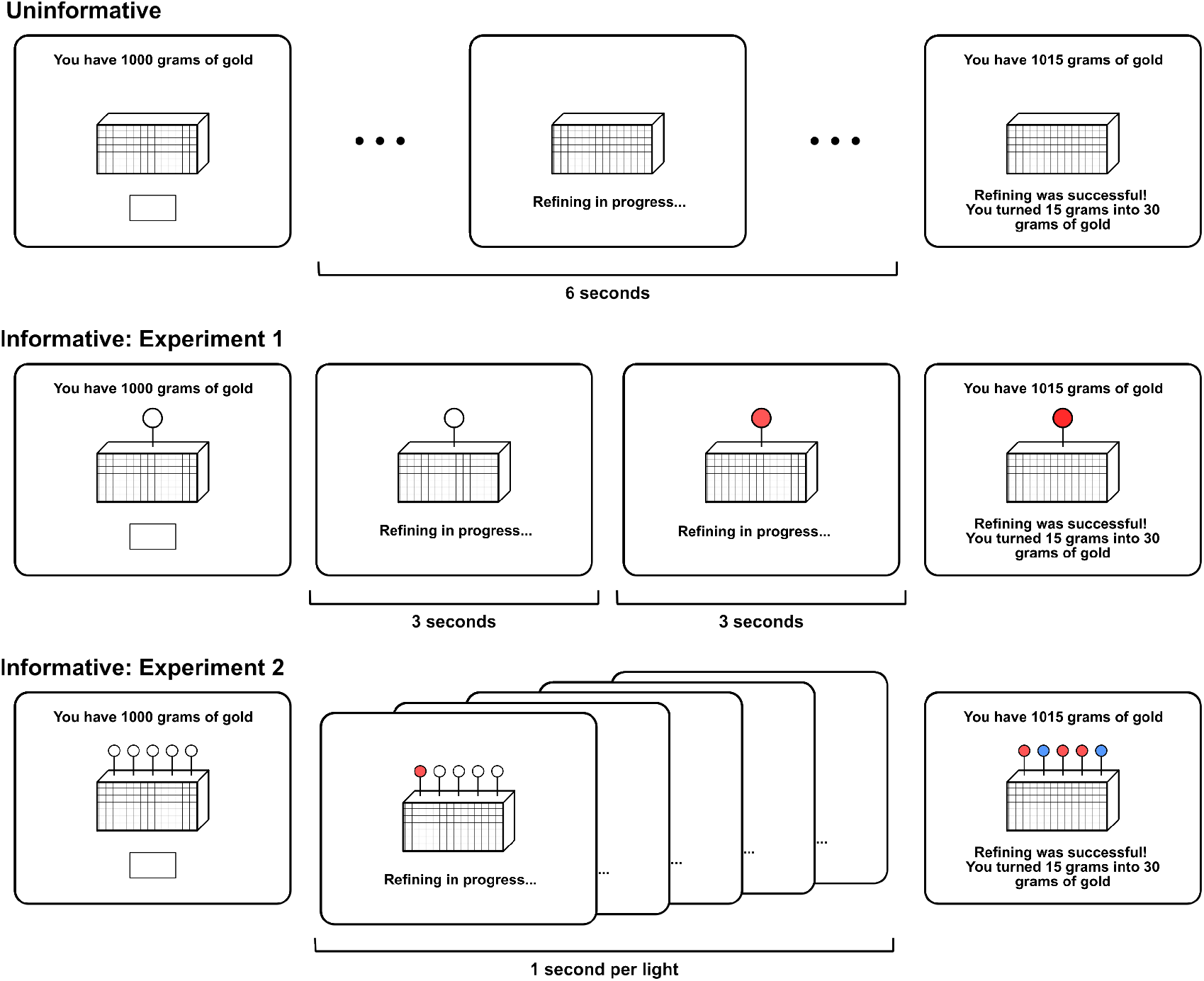
Trial procedure for Experiments 1 and 2. Across all trials, participants were first shown an image of a ‘gold-refining machine’ and were prompted to enter the number of grams of gold they would like to put into the machine in a text box. Each machine was accompanied by text that indicated the probability of successful refinement, the return per gram if refinement was successful, and (for informative machines) the meaning of the different light colours that could be presented (text not shown in this diagram). The machine would process the gold for six seconds, after which it would either return a larger amount of gold than was put in (if successful), or return no gold (if unsuccessful). In uninformative trials, no information was provided about the outcome until the end of the trial. In Experiment 1, informative trials revealed the outcome after three seconds, indicated by the colour change of a light on top of the machine. In Experiment 2, the outcome was revealed by a sequence of five lights switching on with a one-second delay between each.

Importantly, some refinement machines (termed “informative machines”) were accompanied by indicator lights that changed colour during the trial depending on whether the refinement process would be a success or a failure. By contrast, other machines (termed “uninformative machines”) were not. The colours that indicated success and failure were counterbalanced across participants, and participants were explicitly instructed which indicator light colour corresponded to which outcome before each trial. In Experiment 1, informative machines had a single indicator light that changed colour three-seconds into the refinement process, indicating the outcome three seconds earlier than uninformative machines. Importantly, this timing was consistent with the reveal of the third window in the paradigm employed by Matthews et al. (2023), which was the earliest possible time at which the outcome of a trial could be known in their task. This ensured that the indicator light on informative machines in Experiment 1 always resolved uncertainty at least as early as the ‘slot machines’ in the Matthews et al. study. In Experiment 2, we replaced the single light with an array of five lights that changed colour from left to right at one-second intervals. Participants were instructed that each light represented one step of the refinement process, and that the overall success of the refinement was determined by whether a threshold number of refinement steps were successful (indicated by at least three matching lights for trials with a success probability of 50%, and by at least four matching lights for trials with a success probability of 18.75%; see Figure 1).

After completing either task, participants who had completed the PGSI questionnaire were presented with gambling help resources relevant for their self-reported country of residence. Participants with a PGSI score between 5 and 7 were additionally informed that their responses placed them in the “moderate risk” category for gambling-related harm, while those with a score of 8 or above were informed that they fell in the “high risk” category. This process was required by the human research ethics committee that approved the experimental design.

Finally, we note that the first 50 participants in Experiment 1 were exposed to a different type of uninformative trial, in which an uninformative indicator light was shown and changed colour after three seconds, but with no correspondence between the colour change and the refinement outcome. Participants were informed that the light would be uninformative prior to each trial. After collecting the initial sample of 50, we removed the light in the uninformative condition to increase the salience of the contrast between informative and uninformative machines.

### Trial Structure

Each experiment included 76 trials, arranged into four blocks of 19, with self-timed breaks administered between each block. Four trials were attention-checks, in which the participant was instructed that the machine would only refine successfully if they put exactly 16 grams into it, returning 32 grams. The position of these trials was pseudorandomised such that two would be shown within the first two blocks, with the other two in the final two blocks. In both experiments, the order of the experimental trials was fully randomised.

In Experiment 1, each machine had either a 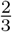 (67% low-risk), 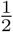 (50%; medium-risk), or 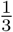 (33%; high-risk), probability of successful refinement. Combined with the two machine types (informative and uninformative), this produced six unique trial types. Each of these trial types was presented 12 times (in a randomised order), for a total of 72 experimental trials. By contrast, in the Matthews et al. (2023) study, all ‘slot machines’ had a 50% probability of reward. The 50% reward condition of Experiment 1 therefore represented the closest conceptual replication of previous work, and the 67% and 33% conditions represented an extension of previous research.

In Experiment 2, each machine required either at least three of the five refinement steps to be successful (50%; medium-risk) or at least four successful steps (18.75%; high-risk). Alongside the two machine types, this produced four unique trial types, each repeated 18 times for a total of 72 experimental trials.

### Data Analysis

All analyses were conducted in R using the *ez* package. To ensure results were robust, we repeated all analyses after excluding participants who failed one or more attention-check trials (N = 18 in Experiment 1, N = 19 in Experiment 2). Excluding these participants had no meaningful effects on the overall results, so we retained all participants for the reported analyses. Similarly, results were unchanged when excluding participants from Experiment 1 who took part in the earlier version of the study (in which uninformative machines had an unpredictive indicator light rather than no light at all).

## Results - Experiment 1

Figure 2A displays mean bet size (amount of gold put into the machine) across our six trial types (3 success probabilities × 2 machine types). The figure displays a large increase in bet size with higher success probability, as well as a more modest increase in bet size for informative over uninformative machines. A 2 × 3 within-subjects ANOVA confirmed significant main effects of both success probability, *F* (2, 298) = 120.95, *p* < .001, generalised *η*^2^ = .151, and machine type, *F* (1, 149) = 5.78, *p* = .017, generalised *η*^2^ < .001. There was no significant interaction between the two factors, *F* (2, 298) = 1.06, *p* = .349. To examine which trial types were producing the significant omnibus main effect of machine type, we conducted post-hoc t-tests with Bonferroni-Holm adjustments for family-wise errors. These tests revealed significant differences between machine type in the 33% success trials (i.e. ‘high-risk trials’), *t* (149) = 2.85, adjusted *p* = .015, but not in the 50% (*t* (149) = 0.63, adjusted *p* = .741), nor the 67% success trials (*t* (149) = 0.90, adjusted *p* = .741).

**Figure 2:**
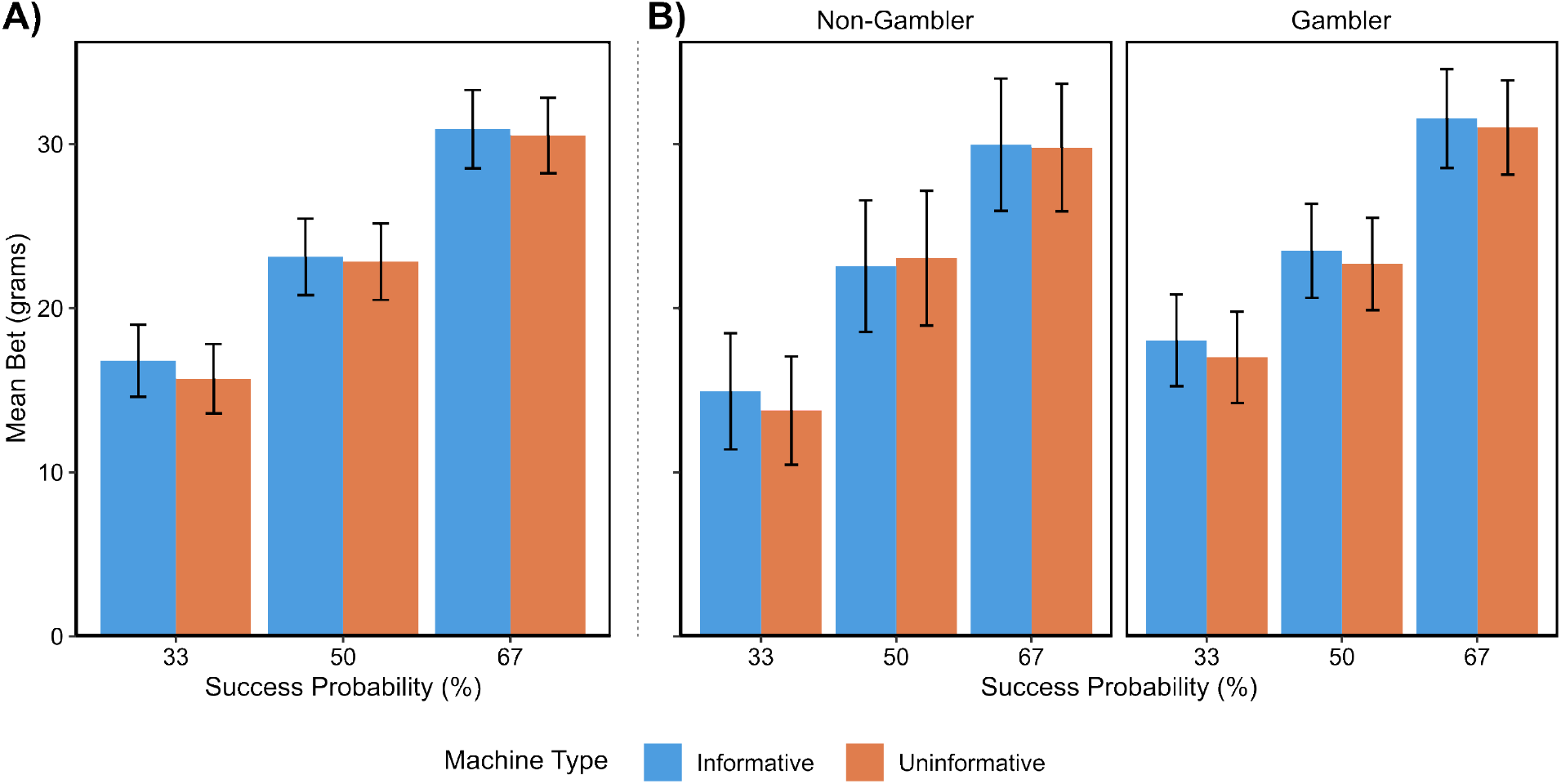
Experiment 1 results. A) Mean bet size across machine type and success probability. B) Mean bet size across trial types split by gambling status (non-gambler: PGSI = 0; gambler: PGSI > 0). All error bars represent 95% confidence intervals.

Next, we set out to test whether gambling status affected bet size (Figure 2B). We first defined two groups: non-gamblers (those who self-reported no engagement with gambling products in the past 12 months, as well as those with PGSI scores of 0; N = 60) and gamblers (those with a PGSI score above 0; N = 90). We next conducted a 2 × 3 × 2 mixed-effects ANOVA, with factors of machine type (2 levels: uninformative or informative), success probability (3 levels: 33%, 50%, or 67%), and gambling status (2 levels: non-gambler or gambler). This analysis reconfirmed the significant main effects of success probability, *F* (2, 296) = 118.99, *p* < .001, generalised *η*^2^ = .149, and machine type, *F* (1, 148) = 4.64, *p* = .033, generalised *η*^2^ < .001. However, we found no significant main effects or interaction effects of gambling status, either at the omnibus level (Supplementary Table 1) or the post-hoc t-tests after Bonferroni-Holm corrections were applied (Supplementary Table 2).

## Interim Discussion

Results of Experiment 1 indicated a strong, positive effect of success probability on bet size. By contrast, our primary research question about the effect of early information on bet size revealed relatively equivocal results. Although there was a significant main effect of machine informativeness on bet size, this effect was substantially smaller than was expected given the magnitude of the effect reported by Matthews et al. (2023). Moreover, exploratory post-hoc analyses of the different machine-risk conditions revealed no evidence for an effect of non-instrumental information within the 50%-win condition that most strongly resembled the ‘slot machine’ gambles utilised by Matthews et al.

In Experiment 1, informative stimuli delivered complete information all at once, revealing the true outcome of the trial after 3 seconds. This represents a point of difference between Experiment 1 and the design of Matthews et al. (2023), which employed a sequential reveal across five windows, with the majority colour determining the final outcome. To determine whether this point of difference may have influenced our findings in Experiment 1, we conducted a second experiment in which our informative machines mimicked this sequential reveal of information. In Experiment 2 we focused on the effects of non-instrumental information in machines with either a 50% (to replicate Matthews et al.) or an 18.75% success probability (determined by the number of successful steps required; see Methods). We included the second condition because Experiment 1 only revealed a significant effect of non-instrumental information in the lowest probability condition (33% success probability). We therefore reasoned that if the effect of non-instrumental information on risk appetite was strongest for low-probability outcomes, a condition with a lower success probability would provide the strongest test of this effect.

## Results - Experiment 2

Figure 3A displays mean bet size across the four trial types (2 success probabilities × 2 machine types). Consistent with Experiment 1, we observed larger bets in the condition with higher success probability. A 2 × 2 within-subjects ANOVA confirmed the significant main effect of success probability, *F* (1, 101) = 72.66, *p* < .001, generalised *η*^2^ = .115. However, we did not observe significant effects of machine type, either in its main effect, *F* (1, 101) = 3.19, *p* = .077, or in its interaction with success probability, *F* (1, 101) = 1.77, *p* = .187. Exploratory post-hoc t-tests with a Bonferroni-Holm correction for multiple comparisons revealed no difference in bet size between informative and uninformative machines for trials with a 18.75% success probability, *t* (101) = 0.47, adjusted *p* = .640. For machines with a 50% success probability, post-hoc analyses found only a marginally significant effect that did not retain significance after correcting for multiple comparisons, *t* (101) = 2.00, adjusted *p* = .097.

**Figure 3:**
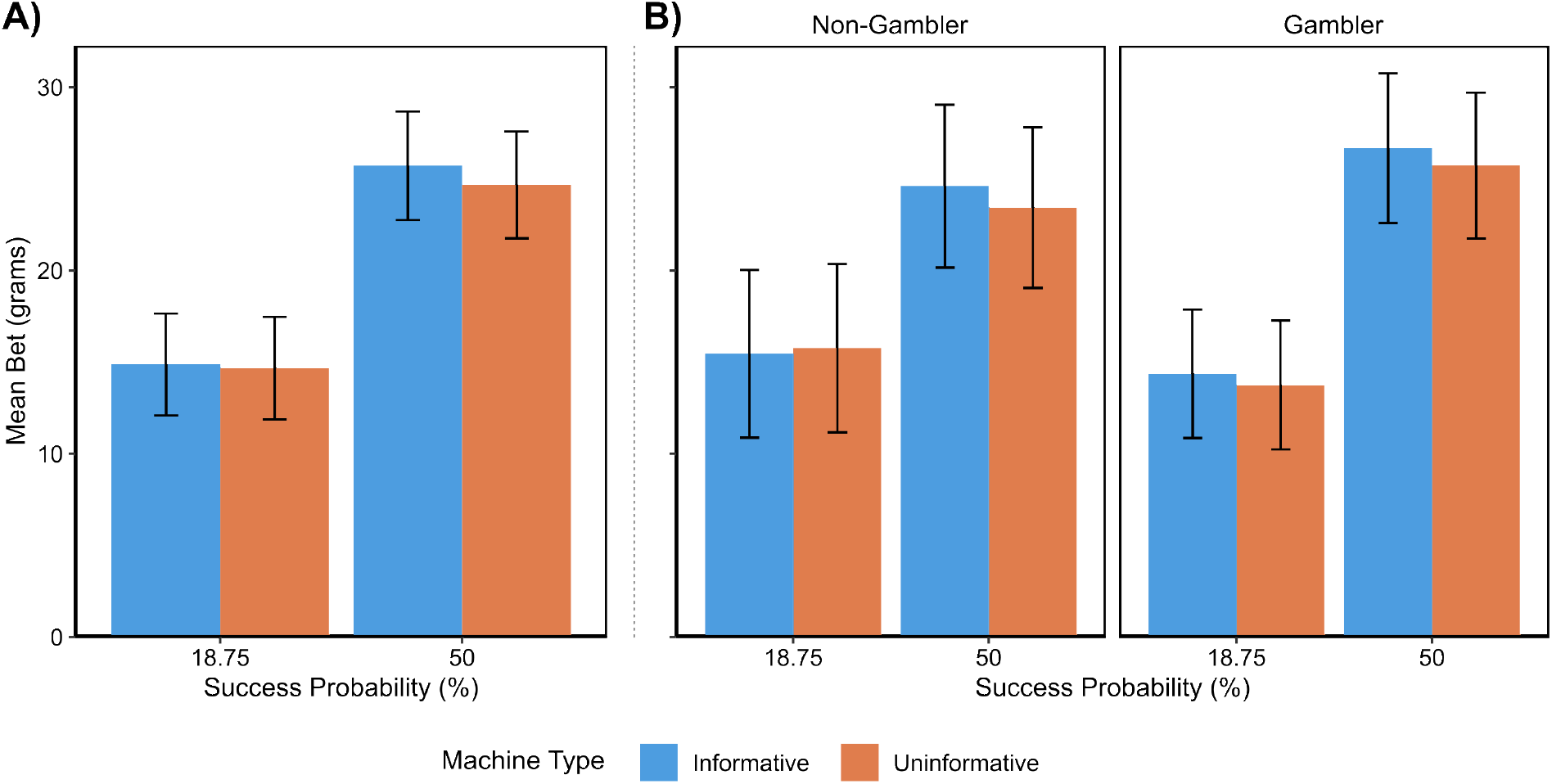
Experiment 2 results. A) Mean bet size across machine type and success probability. B) Mean bet size across trial types split by gambling status (non-gambler: PGSI = 0; gambler: PGSI > 0). All error bars represent 95% confidence intervals.

Finally, we attempted to examine whether these effects differed between the non-gamblers and gamblers in our sample (Figure 3B). We divided our sample into non-gamblers (including those with a PGSI score of 0; N = 47) and gamblers (with a PGSI score above 0; N = 55). A 2 × 2 × 2 mixed-effects ANOVA, with factors of machine type (2 levels: uninformative or informative), success probability (2 levels: 18.75% or 50%), and gambling status (2 levels: non-gambler or gambler) revealed a significant main effect only of success probability, *F* (1,100) = 71.13, *p* < .001, generalised *η*^2^ = .112. We found no other significant main effects or interaction effects in either the omnibus ANOVA (Supplementary Table 3), nor in post-hoc comparisons conducted to examine the effect of machine type in each gambling group at each success probability (Supplementary Table 4).

## Discussion

In this study, we aimed to uncover the cognitive mechanisms that determine the effect of non-instrumental information on decision making under risk. In Experiment 1, we first attempted to replicate the findings of Matthews et al. (2023), that non-instrumental information availability increases risk appetite, using a simplified task in which participants chose how much to bet on the outcome of a fair gamble. Crucially, the outcome of this gamble was either revealed after a six-second delay or signalled halfway through the delay period. We found only weak evidence for an effect of the early reveal of information on risk appetite as measured by participants’ bet amount. We next altered the paradigm in Experiment 2 to use a five-stage sequential reveal of information, thereby increasing the consistency of our task with the original study. However, this adjustment did not strengthen the observed effect of early information on risk appetite: we found no significant overall main effects or interaction effects for non-instrumental information on bet amount. Though the lack of any consistent effects in the present study prevented us from examining the cognitive mechanisms underlying any effect, these findings may nonetheless provide insights into the relationship between non-instrumental information and risk appetite.

One important difference between the current task and that utilised by Matthews et al. is that in our task, participants only determined the size of their wager and were required to participate in the lottery irrespective of that decision. In the original paradigm used by Matthews et al., participants chose whether or not to play the lottery for a fixed bet size. If participants chose not to play, they were required to sit through a sequence of uninformative signals and not told the outcome of the original trial. In our task, participants received information about all outcomes and were shown the iterative display of non-instrumental information on informative trials, even if they placed a very low bet. This was an intentional design decision to decouple preference for observing non-instrumental information from the effects of non-instrumental information on risk appetite.

This difference between task paradigms is likely to explain the difference in findings between the two studies. In particular, we suggest that participants in the study by Matthews et al. may have been more likely to play informative lotteries because they valued the non-instrumental information itself, rather than because the availability of that information increased their risk appetite. This is consistent with previous findings that demonstrate the subjective value of non-instrumental information (Bennett et al., 2016; Charpentier et al., 2018; Goh et al., 2021; Jiwa & Gottlieb, 2026; Van Lieshout et al., 2018). This explanation would also account for the absence of an effect in the present study (or an effect of smaller magnitude, as suggested by some post-hoc results), because information in our informative trials was provided irrespective of what bet size the participant chose.

One alternative explanation could also account for these divergent findings, though we consider it to be less plausible than the informational value hypothesis. It is noteworthy that the measurement of risk appetite differed between studies: whereas Matthews et al. (2023) utilised a binary ‘willingness-to-bet’ metric, the present study measured the magnitude of the bet itself. It is theoretically possible that non-instrumental information lowers the threshold for entering a gamble without necessarily scaling the magnitude of the risk one is willing to take once committed. However, this remains a speculative distinction requiring direct empirical testing, and standard accounts of decision making under risk do not draw a meaningful theoretical distinction between willingness to gamble versus preferred bet size when gambling. In standard versions of prospect theory, for instance, an agent who prefers a gamble over its certainty equivalent for a given wager size and success probability should also strictly prefer larger wagers over smaller wagers for that success probability when choosing a bet amount (and vice versa: those who prefer the certainty equivalent should strictly prefer smaller wagers).

Together, our results suggest that the presentation of non-instrumental information has little-to-no effect on risk appetite as measured by bet size. It is important that we note, however, that this does not preclude the possibility that the presentation of non-instrumental information in gambling products increases the likelihood of gambling-related harm to users through different means. Previous studies have shown that non-instrumental information can be manipulated to encourage misunderstanding of gambling odds or outcomes, such as ‘losses disguised as wins’ (Dixon et al., 2010; Graydon et al., 2021) and ‘near-misses’ (Reid, 1986). Alternatively, it is possible that non-instrumental information may increase user engagement and thereby lead to lower attrition rates over time, or that it may alter the reinforcement-learning processes by which people develop preferences for engaging with gambling products. In each case, further research is required to explore these possibilities.

## Data & Code Availability

All experiment code utilised and data generated and analysed in this study are publicly available in the Open Science Framework: https://osf.io/juy2s.

## Acknowledgements

This work was supported by a Future Fellowship to DB from the Australian Research Council (FT240100555). We are grateful to Trevor Chong for helpful comments on an earlier draft of this report.

